# ApoE4 disrupts interaction of sortilin with fatty acid-binding protein 7 essential to promote lipid signaling

**DOI:** 10.1101/2021.05.20.444938

**Authors:** Antonino Asaro, Rishabhdev Sinha, Magda Bakun, Oleksandra Kalnytska, Anne-Sophie Carlo-Spiewok, Tymon Rubel, Annemieke Rozeboom, Michal Dadlez, Bozena Kaminska, Eleonora Aronica, Anna R. Malik, Thomas E. Willnow

## Abstract

Sortilin is a receptor for neuronal uptake of apolipoprotein E. Sortilin-dependent uptake of lipidated apoE promotes conversion of polyunsaturated fatty acids (PUFA) into neuromodulators that induce anti-inflammatory gene expression in the brain. This neuroprotective pathway works with apoE3 but is lost with apoE4, the main risk factor for Alzheimer’s disease (AD). Here, we elucidated steps in cellular handling of lipids through sortilin, and why they are disrupted by apoE4. Combining unbiased proteome screens with analyses in mouse models, we uncover interaction of sortilin with fatty acid-binding protein (FABP) 7, the intracellular carrier for PUFA in the brain. In the presence of apoE3, sortilin promotes functional expression of FABP7 and its ability to elicit lipid-dependent gene transcription. By contrast, apoE4 binding blocks sortilin sorting, causing catabolism of FABP7 and impairing lipid signaling. Reduced FABP7 levels in the brain of AD patients expressing apoE4 substantiate the relevance of these interactions for neuronal lipid homeostasis. Taken together, we document interaction of sortilin with mediators of extracellular and intracellular lipid transport that provides a mechanistic explanation for loss of a neuroprotective lipid metabolism in AD.

**SUMMARY STATEMENT:** Lipids are central to brain health and defects in brain lipid homeostasis are causal to neurodegenerative processes in Alzheimer’s disease. Here, we uncovered how the neuronal lipoprotein receptor sortilin interacts with apoE and FABP7, the carriers for extra- and intracellular transport of lipids in the brain, respectively. We show that this interaction enables lipids to control gene transcription via nuclear receptors; and why this presumed neuroprotective lipid action is disturbed in humans who carry the ε4 variant of apoE, the most important risk factor for sporadic Alzheimer’s disease.

## INTRODUCTION

Apolipoprotein E is the main carrier for lipids in the brain. It is released by astrocytes and microglia and delivers essential lipids to neurons that take up apoE-bound cargo through apoE receptors expressed on the neuronal cell surface (reviewed in (Holtzman et al., 2012)). ApoE also bears significance as the most important genetic risk factor for the sporadic form of Alzheimer’s disease (AD) as carriers of the *APOEe4* allele are at a significantly higher risk of developing AD than individuals having the common *APOEe3* gene variant (Corder et al., 1993).

The identification of sortilin as a receptor for neuronal clearance of apoE has shed some light on cellular mechanisms implicating this apolipoprotein in brain lipid metabolism and AD progression (Carlo et al., 2013). In detail, sortilin directs neuronal uptake and conversion of apoE-bound ω3-polyunsaturated fatty acids (PUFA) into endocannabinoids (eCBs), neuromodulatory lipids that act via nuclear receptors of the PPAR family to induce an anti-inflammatory gene expression profile in the brain (Asaro et al., 2020). This neuroprotective action of sortilin is seen with apoE3 but lost when binding apoE4, disrupting neuronal eCB production and resulting in a pro-inflammatory state that may predispose the apoE4 brain to neurodegeneration (Asaro et al., 2020). Still, the molecular mechanism of sortilin’s action in neuronal lipid homeostasis and why this activity is lost in the presence of apoE4 remains poorly understood.

Sortilin is a member of the vacuolar protein sorting 10 protein (VPS10P) domain receptor family, a group of specialized receptors involved in endocytosis and intracellular sorting of ligands (reviewed in (Carlo et al., 2014)). Binding of lipid-laden apoE4 to sortilin on the cell surface does not block internalization but disrupts succeeding steps in intracellular receptor sorting (Asaro et al., 2020; Carlo et al., 2013). This observation suggests that it is not the delivery of PUFA into neurons but subsequent steps in intracellular handling of the lipids that are disrupted in the apoE4 brain. Here, we aimed at dissecting the distinct steps of sortilin action in neuronal eCB metabolism impaired by apoE4. Combining unbiased proteome screens in primary neurons and glia with studies in mouse models and AD patient specimens, we uncovered the functional interaction of sortilin with fatty acid-binding protein (FABP) 7, the intracellular carrier for PUFA and eCBs in the brain. ApoE4 disrupts the ability of sortilin to promote stability and proper intracellular sorting of FABP7, essential for lipid signaling via PPARs. These findings identified a central role for sortilin in neuroprotective lipid metabolism by integrating extracellular (apoE) and intracellular (FABP7) lipid transport processes; processes disrupted by apoE4-induced missorting of this receptor.

## RESULTS

### Surface proteome analyses identify FABP7 as novel sortilin target in neurons

Sortilin is a sorting receptor that directs proteins between the cell surface and various intracellular compartments. To identify sortilin-dependent trafficking processes relevant for neuronal lipid metabolism and action, we applied an unbiased proteomics approach to identify novel receptor targets in neurons. This approach was based on our assumption that the loss of sortilin activity will result in aberrant distribution of so far unknown receptor ligands between cell surface and intracellular compartments. Such unbiased screens had been used successfully by us before to identify targets for related VPS10P domain receptors (Malik et al., 2019; Subkhangulova et al., 2018).

To identify sortilin ligands, we biotinylated surface proteins in primary neurons from wild-type mice (WT) or animals genetically deficient for *Sort1*, the gene encoding sortilin (KO) (Jansen et al., 2007). The biotinylated proteins were purified from cell extracts using streptavidin beads, identified by LC-MS/MS, and subjected to LC-MS label-free quantitation (workflow in Fig. 1A). By this approach, we identified multiple proteins with altered abundance in the neuronal cell surface fraction upon loss of sortilin (Fig. 1B); some of which may be primary, others secondary targets of receptor dysfunction in KO neurons. Proteins with altered abundance in the cell surface fraction of KO neurons included known receptor ligands, such as tropomyosin receptor kinase B (TrkB) (Vaegter et al., 2011), EGF receptor (Al-Akhrass et al., 2017), sphingomyelin phosphodiesterase (Ni and Morales, 2006), and the vacuolar protein sorting-associated protein 26B (Kim et al., 2010) (Fig. 1B, Suppl. table 1). These findings corroborated our strategy for identifying primary targets of sortilin-mediated protein sorting. Among the proteins with altered surface abundance in KO neurons was one hit with particular relevance to PUFA and eCB metabolism, namely brain fatty acid binding protein (FABP) 7. The abundance of FABP7 was increased in the surface proteome of KO neurons, suggesting altered subcellular distribution of the carrier in the absence of sortilin (Fig. 1B, Suppl. table 1). This effect was specific to FABP7 and not seen for other fatty acid-binding proteins expressed in the brain, namely FABP3 and FABP5 (Suppl. table 1).

**Figure 1.**
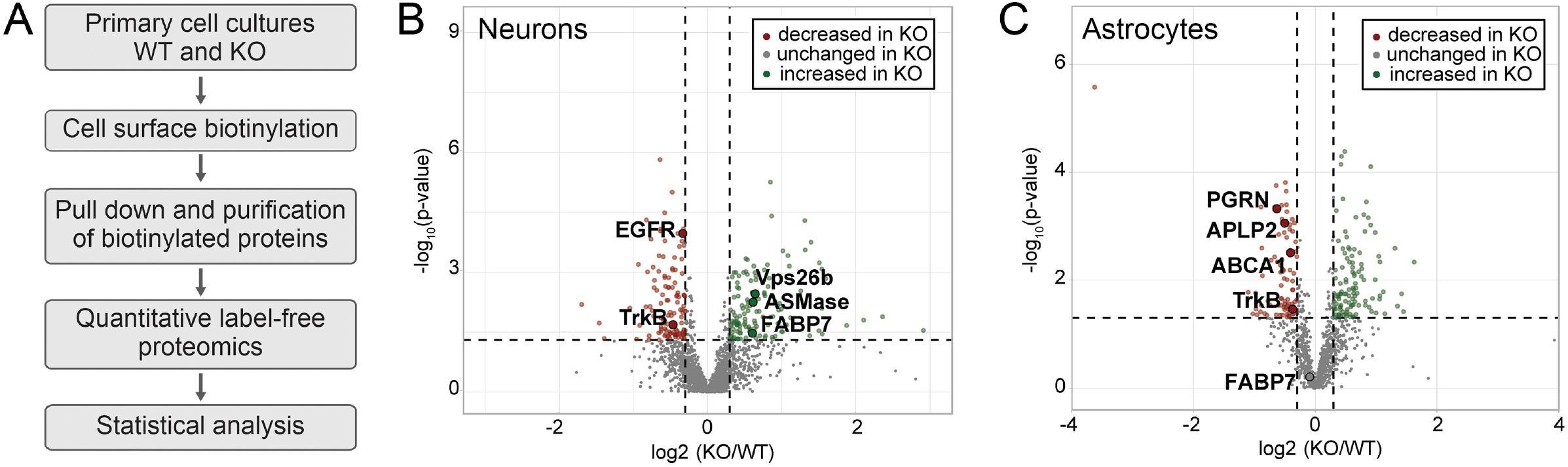
Surface proteome analysis identifies fatty acid-binding protein 7 as novel sortilin target in neurons but not astrocytes. (**A**) Workflow of surface proteome analysis in primary neurons from wild-type (WT) and *Sort1*^-/-^ (KO) mice. (**B**) Comparison of the surface proteomes of WT and KO primary neurons (*n*=3 biological replicates/group; 2 technical replicates per biological replicate). Dashed lines show threshold values for log2 (fold change) and −log10 (*p*-value), ± 0.3 and 1.3, respectively. Proteins with increased (green) or decreased (red) levels in the surface proteome of KO neurons are color-coded. Selected proteins with altered surface exposure in KO neurons are highlighted (see Suppl. table 1 for details). (**C**) Data as in panel B but comparing the surface proteomes of WT and KO primary astrocytes (*n*=6 biological replicates/group). Selected proteins with altered surface exposure in KO astrocytes are highlighted (see Suppl. table 1 for details).

To query the specificity of sortilin-dependent sorting of FABP7 in neurons, we repeated the quantitative surface proteome analysis in primary astrocytes from WT and KO mice (Fig. 1C). Again, established sortilin interaction partners were identified as being altered in their presence in the cell surface fraction, including TrkB, amyloid-like protein 2 (Butkinaree et al., 2015), ATP-binding cassette transporter ABCA1 (Lv et al., 2019), and progranulin (Hu et al., 2010). By contrast, cell surface localization of FABP7 was not impacted by sortilin deficiency in astrocytes (Fig. 1C, Suppl. table 1).

### Sortilin facilitates expression of FABP7 in CHO cells

FABP7 is a brain-specific lipid chaperone that enables cellular uptake and cytosolic trafficking of PUFA and eCBs (Feng et al., 1994; Kaczocha et al., 2009; Kurtz et al., 1994). FABP7-dependent trafficking is required for proper biosynthesis of eCBs from PUFA, and for their delivery to nuclear PPARs for regulation of gene transcription (reviewed in (Moulle et al., 2012)). When myc-tagged FABP7 and sortilin were co-expressed in Chinese hamster ovary cells (CHO-S/F), sortilin co-immunoprecipitated with an affinity resin directed against the myc epitope in FABP7, confirming FABP7 as sortilin ligand (Fig. 2A). Coimmunoprecipitation was also seen in the presence of apoE3 or apoE4, arguing that FABP7 and apoE simultaneously binding to their receptor sortilin (Fig. 2B). Interaction of sortilin with FABP7 was supported by proximity ligation assay (PLA), documenting close proximity of both proteins in intracellular compartments of CHO-S/F cells (Fig. 2C). In subcellular fractionation experiments, FABP7 and sortilin mainly co-localized in the trans-Golgi network (TGN), early endosomes, and the cell surface fraction of CHO-S/F cells (Fig. 2D), consistent with a role for sortilin in sorting of cargo between plasma membrane and secretory and endocytic compartments (Nielsen et al., 2001).

**Figure 2.**
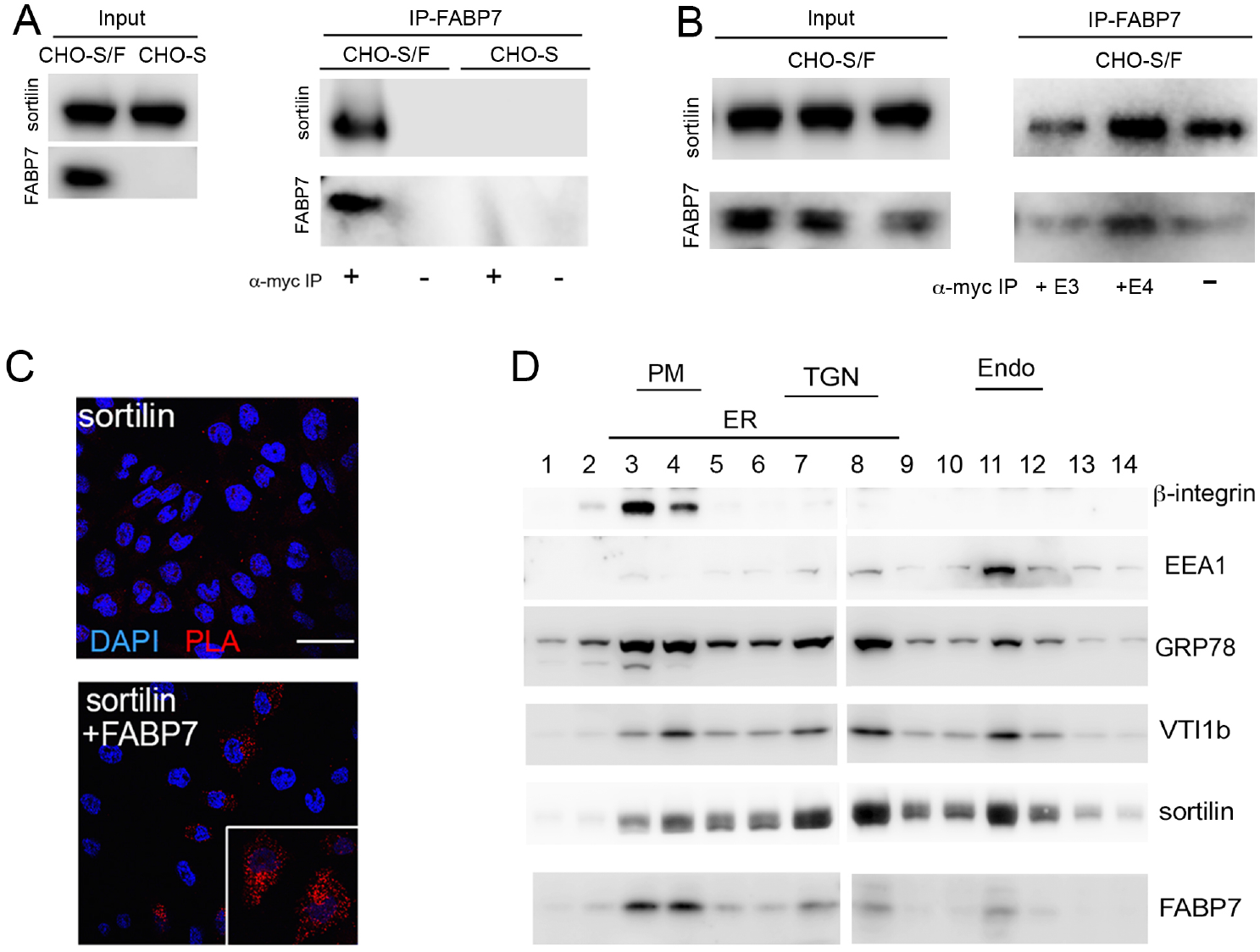
Sortilin and FABP7 interact in cells. **(A**) Chinese hamster ovary (CHO) cells stably overexpressing sortilin (CHO-S) or sortilin and FABP7 tagged with a myc epitope (CHO-S/F) were used for co-immunoprecipitation experiments. Expression of sortilin and FABP7-myc in total cell lysates of both cell lines is shown in the panel Input. In panel IP-FABP7, co-immunoprecipitation of sortilin with anti-myc affinity resin (+) is seen in lysates from CHO-S/F but not from CHO-S cells. No immunoprecipitation of sortilin with anti-myc affinity resin is seen in CHO-S cells, or in CHO-S/F in the absence of the affinity resin (-). (**B**) Co-immunoprecipitation of sortilin with FABP7-myc from CHO-S/F cells as described in panel A. Prior to immunoprecipitation with anti-myc affinity resin, cells were treated with conditioned medium from HEK293 cells containing 5 μg/ml of apoE3 (+E3) or apoE4 (+E4; see Suppl. methods for details) or blank medium (-) for 24 hours. **(C)** Proximity ligation assay (PLA) to assess close spatial proximity of sortilin and FABP7 in CHO cells. Primary antibodies were directed against sortilin or the myc epitope in FABP7-myc. Close proximity is detected in CHO cells expressing both sortilin and FABP7 (red signal). No PLA signal is seen in cells expressing sortilin only. Cell nuclei were counterstained with DAPI (blue). The inset shows a higher magnification image of cells positive for PLA signals for sortilin and FABP7-myc. Scale bar: 50 μm. (**D**) Subcellular fractionation of CHO-S/F cells using gradient ultracentrifugation. Fractions were identified based on markers for endoplasmic reticulum (ER; GRP78), plasma membrane (PM; β-integrin), *trans*-Golgi network (TGN; VTI1b), and early endosomes (Endo; EEA1). FABP7 co-localizes with sortilin to the PM (fractions 3-4), TGN (fractions 7-8), and endosomes (fraction 11).

Of note, levels of FABP7 were significantly higher in CHO cells expressing sortilin (CHO-S) as compared with parental CHO cells lacking the receptor (Fig. 3A-B). This effect was specific for FABP7 as levels of the co-transfected green fluorescent protein (GFP) were identical in both cell lines (Fig. 3A). Elevated levels of FABP7 in CHO-S as compared with parental CHO cells were due to an increase in FABP7 half-life as shown by comparing the turnover of the protein in CHO and CHO-S cells following treatment with cycloheximide, an inhibitor of protein translation (Fig. 3C-D).

**Figure 3.**
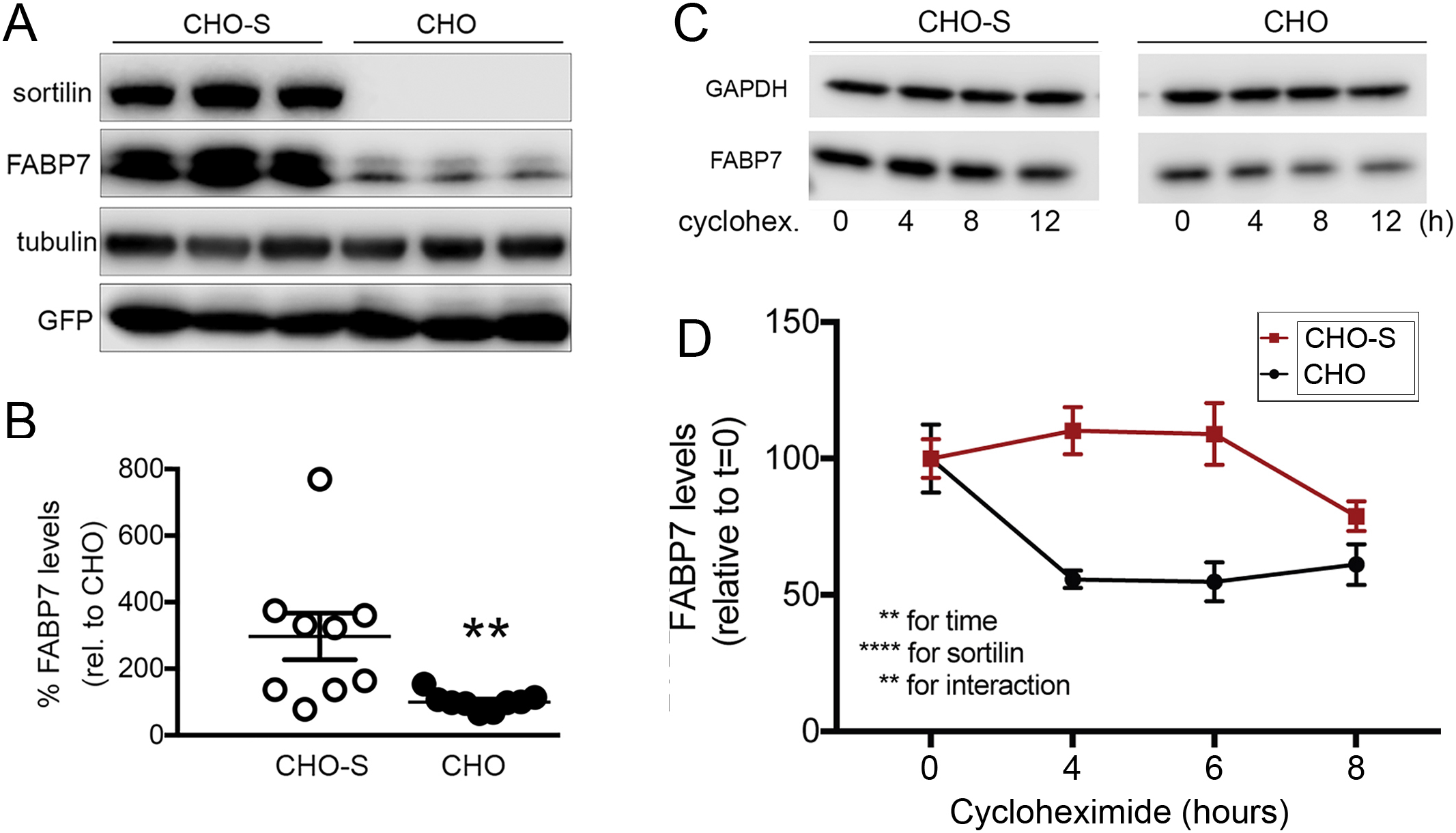
Sortilin stabilizes cellular levels of FABP7 in CHO cells. **(A, B)** Parental Chinese hamster ovary (CHO) cells or CHO cells stably expressing sortilin (CHO-S) were transiently transfected with expression constructs encoding for FABP7 and green fluorescent protein (GFP). An exemplary western blot documenting expression of sortilin and FABP7 in replicate lysates of CHO and CHO-S cells is shown in panel A. Detection of GFP and tubulin served as transfection and loading controls, respectively. Panel B shows FABP7 levels in CHO and CHO-S transfectants as determined by densitometric scanning of replicate western blots (n= 9 replicates from 3 independent experiments per cell line). Data are mean ± SEM given as percent of FABP7 levels in CHO (set to 100%). Levels of FABP7 (but not of GFP) are significantly increased by the presence of sortilin in CHO-S compared with CHO cells. **, p<0.01 (Student’s *t* test). **(C, D)** CHO and CHO-S cells were transiently transfected with a FABP7-myc expression construct. Forty-eight hours post transfection, replicate cultures of transfected cells were treated with 10 μg/ml of cycloheximide and collected at time points 0, 4, 8, and 12 hours later. Levels of FABP7 were determined by western blotting (C). Detection of GAPDH served as loading control. The decrease in FABP7 levels was significantly faster in CHO than in CHO-S cells as determined by densitometric scanning of replicate blots (D; n= 9 replicates per condition from 3 independent experiments). Data are mean ± SEM given as percent of FABP7 levels in CHO or CHO-S at 0 hour of treatment (set to 100%). The significance of data was determined by Two-way ANOVA, followed by Bonferroni post-hoc analysis (**, p<0.01; ****, p<0.0001).

### Sortilin promotes neuronal expression of FABP7 in an apoE isoform-dependent manner

So far, our studies documented a direct interaction of sortilin with FABP7 that controlled stability and intracellular distribution of the lipid carrier in CHO cells (Fig. 3) and neurons (Fig. 1B). To test whether this interaction may also be relevant for control of FABP7 levels *in vivo*, we studied levels of the carrier in brains of WT and KO mice. Given the importance of apoE genotype for sortilin action in brain lipid homeostasis (Asaro et al., 2020), WT and KO mice carried a targeted replacement of the murine *Apoe* locus with genes encoding human apoE3 (E3) or apoE4 (E4) (Knouff et al., 1999). In the presence of apoE3, total brain levels of FABP7 were significantly higher in WT mice as compared to animals lacking sortilin (E3/WT versus E3/KO; Fig. 4A and C). However, in apoE4 mice, sortilin did not impact FABP7 expression as levels of the carrier were comparable in E4/WT and E4/KO animals (Fig. 4B and C). As a control, sortilin deficiency did not affect levels of FABP5 in E3 or E4 animals (Fig. 4A-B). The beneficial effect of sortilin activity on FABP7 levels in E3 mice manifested through a post-transcriptional mechanism as brain levels of *Fabp7* transcripts were not altered comparing E3 and E4 animals, either WT or KO for *Sort1* (Fig. 4D). These experiments substantiated the importance of sortilin for promoting FABP7 levels in the brain of mice expressing apoE3, but not apoE4. An effect of apoE isoform on FABP7 levels was also seen in the human AD brain as levels of the protein were significantly higher in patients of the *APOEε3/APOEε3* as compared with the *APOEε4/APOEε4* genotype (Fig. 4E-F, Suppl. table 2). As in the mouse brain, levels of FABP5 were comparable in both genotypes (Fig. 4E).

**Figure 4.**
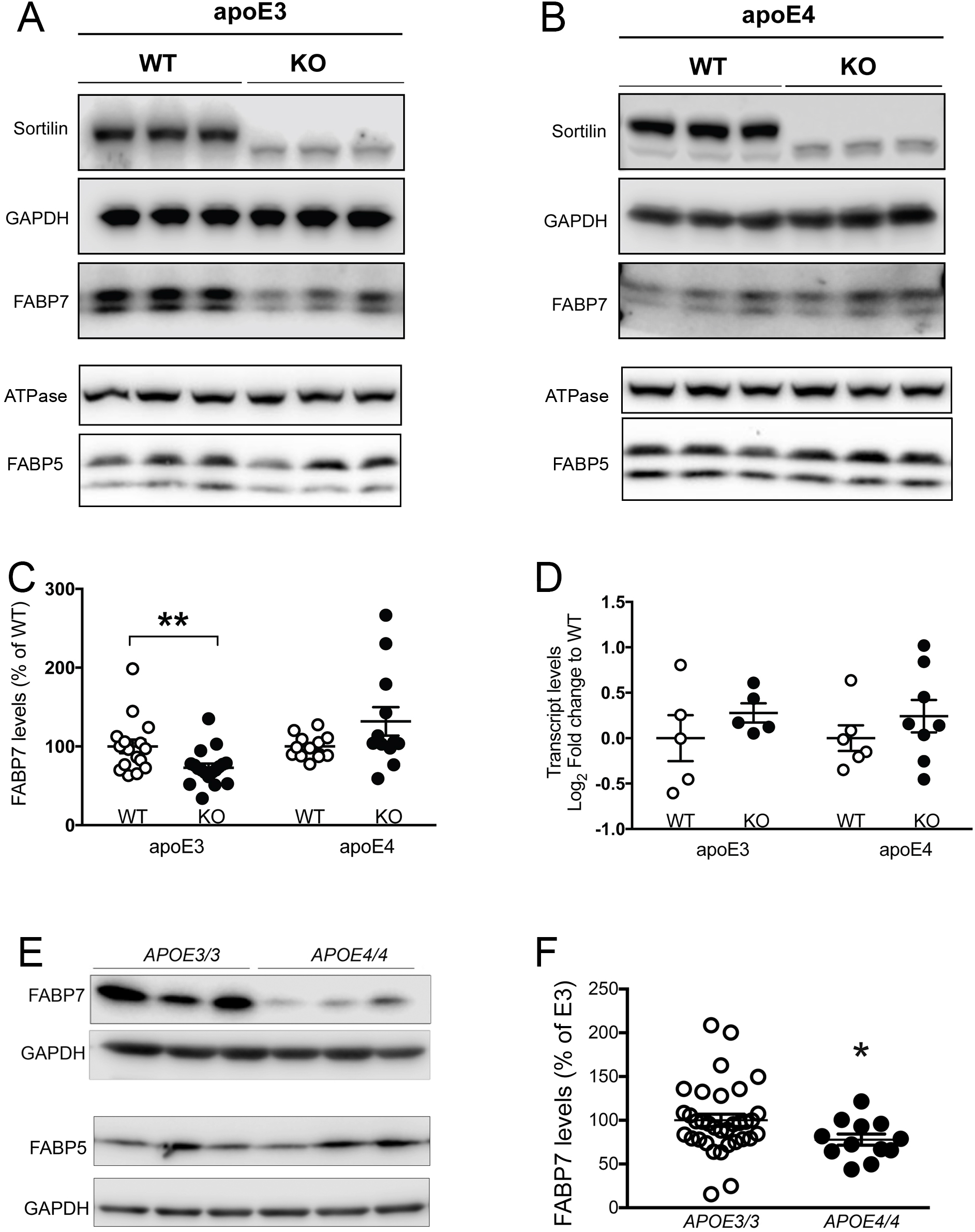
Sortilin deficiency decreases FABP7 levels in brains of apoE3 but not apoE4 mice. **(A, B)** Western blot analysis of FABP5 and FABP7 levels in brain cortices of apoE3 (A) and apoE4 (B) targeted replacement mice, either wild-type (WT) or homozygous for the *Sort1* null allele (KO) (at 3 months of age). Na/K ATPase and GAPDH served as loading controls for detection of FABP5 and FABP7, respectively. Detection of sortilin served as genotype control. (**C**) Quantitative analysis of FABP7 levels in brain cortices of apoE3- and apoE4-expressing mice of the indicated *Sort1* genotype using densitometric scanning of replicate western blots. Values are mean ± SEM given as percent of the respective WT control (set to 100%). Student *t* test was used to determine significance of differences between genotypes (n=12-18 animals per group; **, p<0.01). (**D**) Levels of *Fabp7* transcripts in brain extracts (cortex and hippocampus) of apoE3- and apoE4-expressing mice of the indicated *Sort1* genotypes as determined by quantitative RT-PCR. Values are given as mean ± SEM. No statistically significant differences in transcript levels were observed using Student’s *t* test (n=5-8 animals per group). (**E, F**) Western blot analysis of FABP5 and FABP7 levels in prefrontal cortex specimens of AD patients homozygous for *APOEe3* or *APOEe4* (pathological characteristics given in Supp. table 2). An exemplary western blot is shown in panel E. Detection of GAPDH served as loading control. Panel F shows the result of densitometric scanning of replicate blots. Values are mean ± SEM given as percent of *APOEe3/3* genotype (mean value set to 100%). Welch’s *t*-test was used to calculate the significance of data (n=12-34 individuals per group; *, p<0.05).

In the adult mouse brain, predominant expression of FABP7 has been localized to several types of glia, such as ependyma and Bergmann glia (Kurtz et al., 1994). Still, using immunohistology we also detected moderate but consistent co-expression of FABP7 with sortilin in cortical neurons marked by NeuN (Fig. 5A). Neuronal expression of FABP7 was substantiated in human AD brain tissue (Fig. 5B, Suppl. table 3). To document that decreased levels of FABP7 seen in apoE3 mice upon deletion of sortilin (E3/KO) represented loss from neurons normally co-expressing both proteins, we performed comparative expression analyses in primary neurons and astrocytes from all four genotypes. As in the brain, co-expression of FABP7 with sortilin was seen in cultured WT neurons (Fig. 6A). These cell cultures produced apoE3 and apoE4, likely from astrocytes present in these neuron-enriched cultures (Fig. 6B). The relevance of sortilin and apoE3 for control of neuronal levels of FABP7 was substantiated by western blot analysis documenting significantly higher levels of FABP7 in E3/WT as compared to E3/KO neurons (Fig. 6C-D). By contrast, the presence of sortilin did not affect FABP7 levels in neuronal cultures from E4 mice, in line with a presumed functional deficiency of this receptor in the presence of apoE4 (Fig. 6C-D). As in total brain, loss of sortilin did not impact levels of FABP5 in either apoE genotype (Fig. 6C). The difference in FABP7 levels comparing E3/WT and E3/KO cell preparations was not due to a difference in the amounts of FABP7-producing astrocytes in the cultures, as transcript levels of the astrocyte marker *Gfap* were comparable in both cell preparations (Fig. 6E). Also, levels of *Fabp7* transcript in primary neurons were not impacted by *APOE* or *Sort1* genotypes, substantiating a post-transcriptional mechanism of expression control (Fig. 6F). Robust expression of FABP7 was detected in cultured primary astrocytes from E3/WT and E3/KO mice (Suppl. figure 1A). However, contrary to the situation in primary neurons, FABP7 levels were not altered in primary astrocytes from E3 mice by the presence or absence of sortilin (Suppl. figure 1B-D), nor were the levels of *Fabp7* transcripts (Suppl. figure 1E).

**Figure 5:**
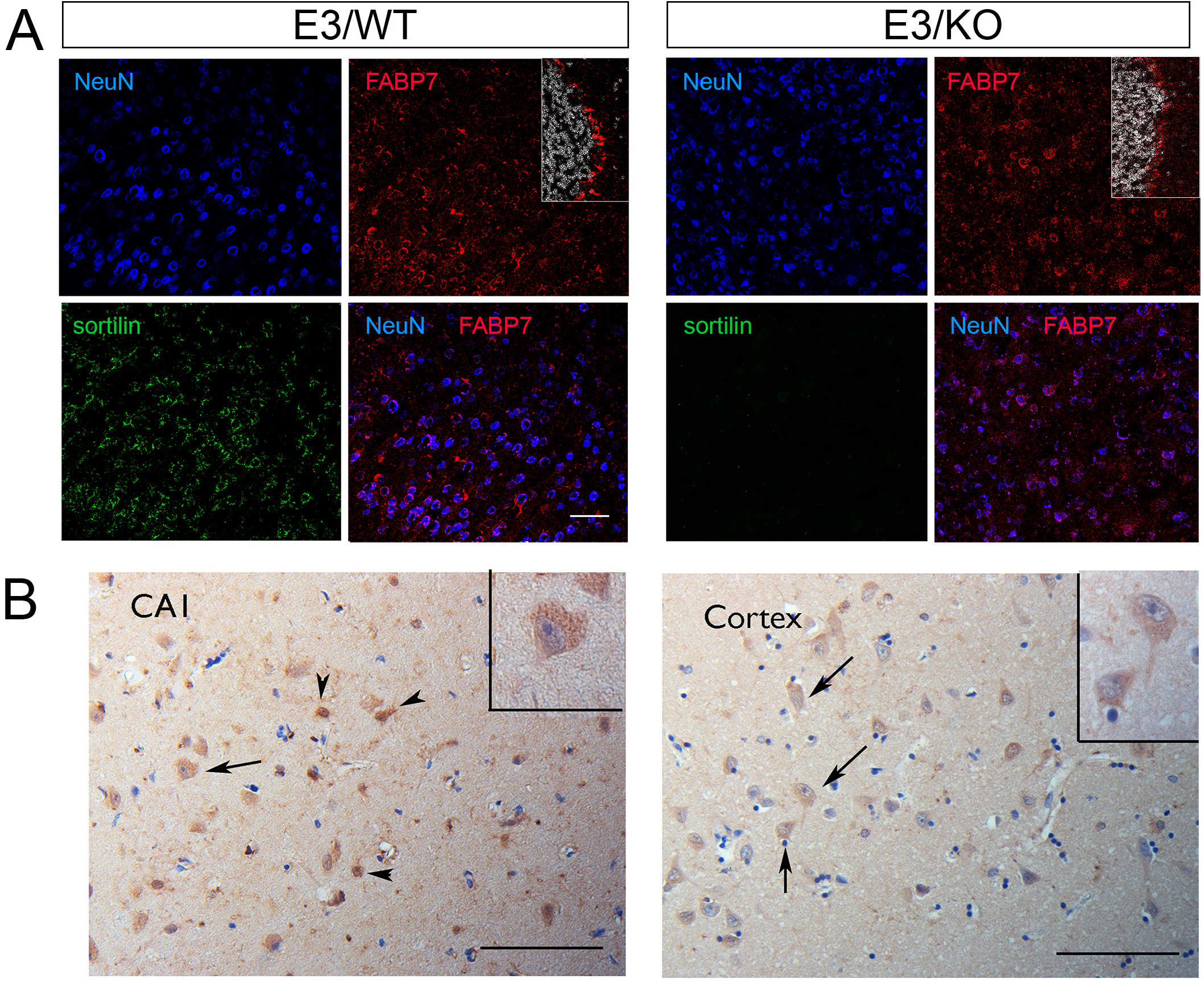
Expression of FABP7 in neurons of mouse and human brains. (**A**) Immunohistological detection of FABP7 (red) in brain cortical sections from apoE3 mice, either wild-type (WT) or genetically deficient for *Sort1* (KO). Additionally, the sections were stained for the neuronal marker NeuN (blue) and sortilin (green) as well as DAPI (white; in insets). Merged images show co-expression of FABP7 with NeuN. As a positive control, the insets document expression of FABP7 in glia in the cerebellum of E3/WT and E3/KO mice. Representative images from analysis of 3 mice per genotype are shown. Scale bar:100 μm. (**B**) Staining for FABP7 in hippocampal subfield CA1 and temporal cortex specimens of AD patients, showing immunoreactivity in glial cells (arrowheads) and light positivity in sparse neuronal cells (arrows, and insets). Representative images from one of three AD cases analyzed are shown (pathological characteristics given in Supp. table 3). Scale bars: 100 μm and 50 μm (inset).

**Figure 6.**
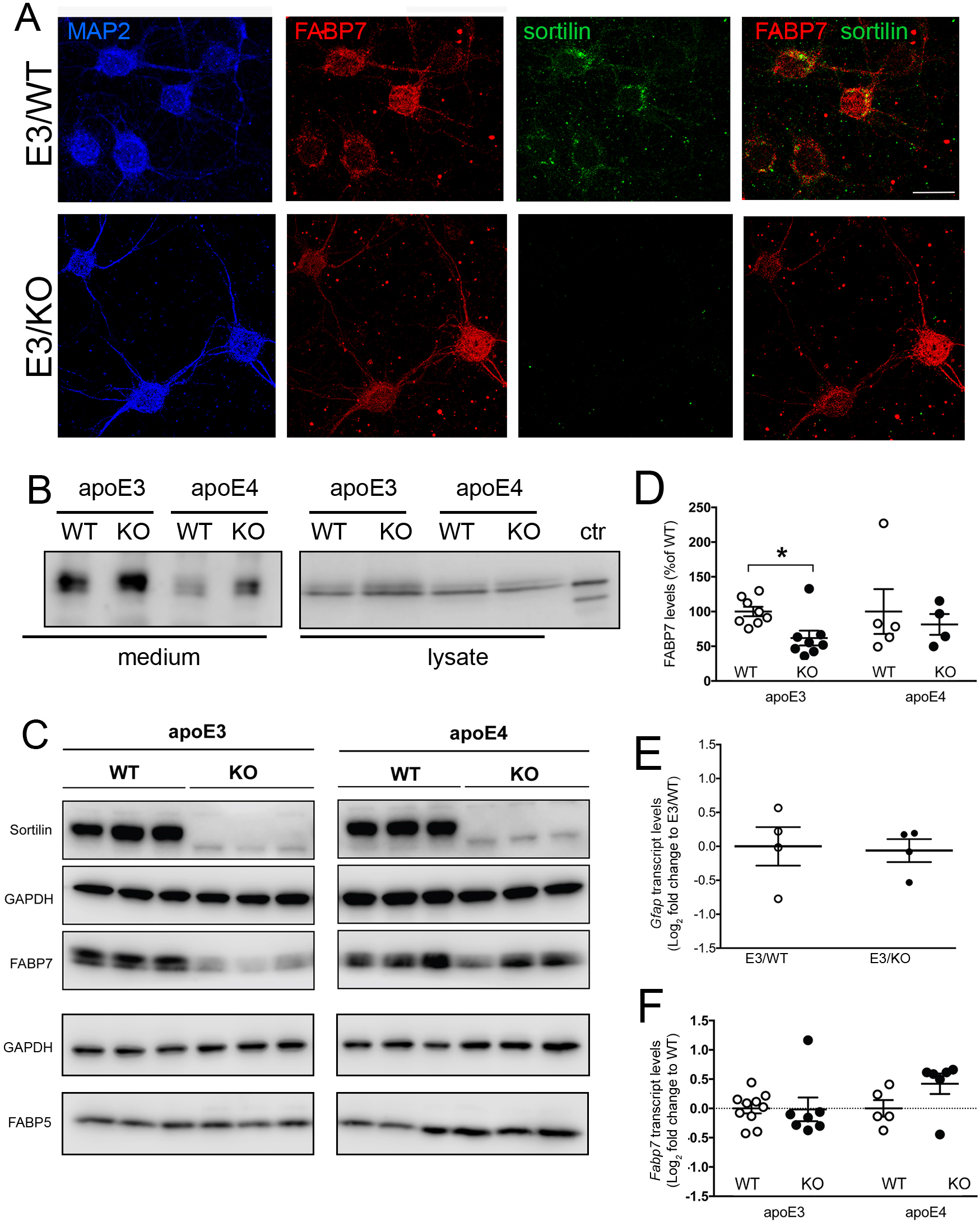
Sortilin deficiency decreases FABP7 levels in primary neurons of apoE3 but not apoE4 mice. **(A)** Immunodetection of FABP7 (red) and sortilin (green) in primary neuronal cultures from apoE3 mice either (WT) or genetically deficient for *Sort1* (KO). FAPB7-stained cells are identified as neurons by expression of MAP2 (blue). Merged images show co-expression of FABP7 and sortilin. Scale bar: 20 μm. (**B**) Detection of apoE3 and apoE4 in medium and lysate of neuronal cultures of the indicated genotypes. Detection of recombinant apoE4 served as control (ctr). **(C-D)** Western blot analysis of sortilin, FABP7, and FABP5 in primary neurons from apoE3 or apoE4-expressing mice, either wild-type (WT) or genetically deficient for *Sort1* (KO). Detection of sortilin and GAPDH served as controls. Levels of FABP7 are reduced in primary cortical neurons from apoE3 mice lacking sortilin (E3/KO) compared with neurons from (E3/WT) animals as determined by densitometric scanning of replicate blots (exemplified in panel C). No statistically significant difference in FABP7 levels is seen comparing neurons from apoE4 mice with (E4/WT) and without (E4/KO) sortilin (panel D). Data are given as mean ± SEM with the respective mean WT levels set to 100% (n=4-8 biological replicates per genotype; Student’s t test). *, p<0.05. (**E**) Quantitative RT-PCR analysis of *Gfap* transcripts in primary neuronal cultures from E3/WT and E3/KO mice. Values are given as log2 fold changes relative to E3/WT set to value 0 (n=4 independent cultures per genotype). (**F**) Levels of *Fabp7* transcript were determined in mouse primary neuronal cultures of the indicated genotypes by quantitative RT-PCR (n=7-10 biological replicates per group for apoE3; n=5-6 biological replicates per group for apoE4). Values are given as log2 fold change relative to transcript levels in the respective WT (set to value 0).

### ApoE4 disrupts the ability of sortilin to control functional expression of FABP7

So far, our findings uncovered a role for sortilin in controlling neuronal stability and subcellular localization of FABP7. The ability of sortilin to sustain FABP7 levels in the murine brain, and in cultured neurons thereof, was seen in the presence of apoE3 but not with apoE4. To explore the underlying molecular mechanisms, we applied the CHO-S/F cell model to study the functional interaction of sortilin with FABP7 in the presence of apoE3 or apoE4 (Fig. 7). As in the murine brain, the ability of sortilin to promote FABP7 levels in CHO cells was seen with apoE3 but compromised by the presence of apoE4. This effect was documented by treating CHO-S/F cells with conditioned medium from HEK293 cells secreting human apoE3 or apoE4 (Chen et al., 2010). Applying identical amounts of both apoE isoforms to the cell medium, levels of FABP7 in CHO-S/F cells were significantly lowered by the addition of apoE4 as compared with apoE3 (Fig. 7A-B). The same effect was seen using native apoE3 or apoE4 secreted by primary astrocytes (Fig. 7C-D).

**Figure 7.**
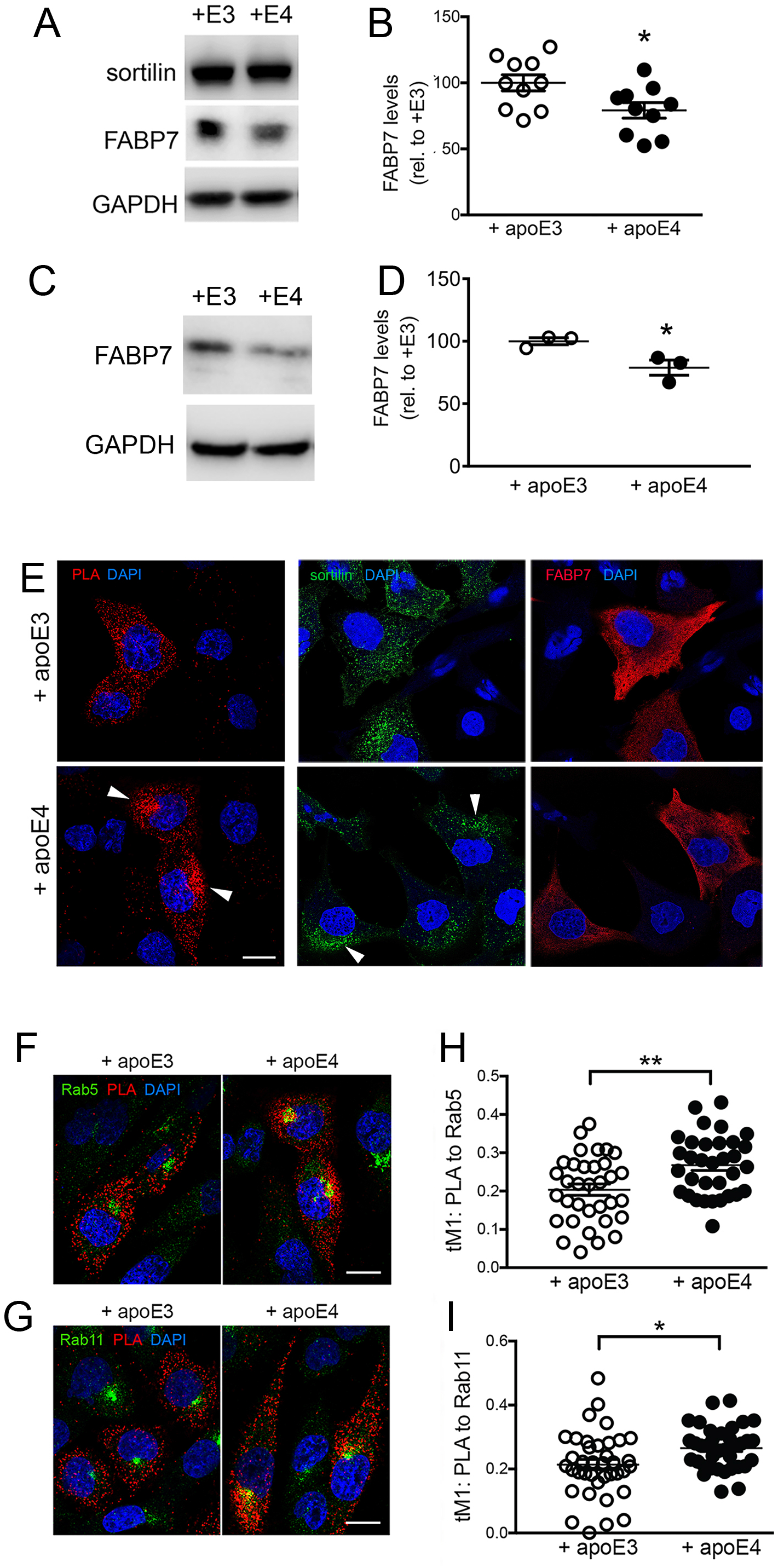
Levels and intracellular sorting of FABP7 are impacted by application of apoE4. (**A-B**) CHO cells stably co-expressing sortilin and FABP7 (CHO-S/F) were treated for 24 hours with conditioned medium containing 5 μg/ml of apoE3 (+E3) or apoE4 (+E4; see methods for details). Western blot analysis (panel A) and densitometric scanning of replicate blots (panel B) document reduced levels of FABP7 in CHO-S/F cells in the presence of apoE4 as compared with apoE3 (n=10 replicate cultures from 4 independent experiments). Data are mean ± SEM given as percent of FABP7 levels in apoE3-treated cells (set to 100%). *, p<0.05 (Student’s *t* test). (**C-D**) Experiment as in panels A and B but using conditioned medium from primary astrocytes secreting apoE3 or apoE4 (n=3 replicates from one culture). Data are mean ± SEM given as percent of FABP7 levels in apoE3-treated cells (set to 100%). *, p<0.05 (Student’s *t* test). (**E**) Proximity ligation assay (PLA) to visualize the intracellular localization of sortilin-FABP7 complexes (red signal) in CHO-S/F cells treated with apoE3- or apoE4-conditioned HEK293 medium for 24 hours (left panel). For comparison, immunostaining of total sortilin (middle panels) and FABP7 (right panels) in treated cells are shown as well. Cell nuclei were counterstained with DAPI (blue). PLA signals for sortilin-FABP7 complexes change from a dispersed vesicular pattern with apoE3 to a perinuclear pattern with apoE4 (arrowheads). A similar change is seen for total sortilin, whereas the pattern for total FABP7 remains unaffected by apoE4. (**F-G**) Colocalization studies in CHO-S/F cells documenting the presence of sortilin-FABP7 complexes (as deduced by PLA; red signal) in early endosomes, marked by antibodies against Rab5 (green signal; panel F) or recycling endosomes marked by Rab11 (green signal, panel G). (**H-I**) Extent of colocalization of PLA for sortilin and FABP7 with Rab5 (panel H) and Rab11 (panel I) as documented by thresholded Mander’s coefficient tM1. This experiment was replicated 3 times (n=33-38 cells per condition). *, *p*<0.05; **, *p*<0.01 (Student’s *t* test). Scale bars: 20 μm

Because apoE4 disrupts intracellular routing of sortilin (Asaro et al., 2020), we tested whether the ability of the receptor to direct sorting of FABP7 may be compromised by binding of apoE4. To verify this notion, we compared the subcellular distribution of sortilin-FABP7 complexes in CHO-S/F cells treated with apoE3 or apoE4. As shown in Fig. 7E, the intracellular distribution of sortilin-FABP7 complexes (as detected by PLA) changed from a dispersed vesicular pattern with apoE3 to a confined perinuclear appearance with apoE4. A similar effect of apoE4 was seen on sortilin alone but not on FABP7 alone (Fig. 7E). These findings corroborate that apoE4 primarily acts on sortilin, impacting sorting of the fraction of FABP7 bound by this receptor. Altered sorting of sortilin-FABP7 complexes in the presence of apoE4 was confirmed quantitatively by colocalization studies documenting trapping of PLA signals for sortilin and FABP7 in early endosomes (marked by Rab5; Fig. 7F and H) and recycling endosomes (marked by Rab11; Fig. 7G and I) in the presence of apoE4 as compared with apoE3.

Among other functions, FABP7 directs PUFA and eCBs to the nucleus to activate transcription factors of the PPAR family (Adida and Spener, 2006; Mita et al., 2010; Tripathi et al., 2017). To explore whether apoE4-induced missorting of sortilin disrupts this FABP7 function, we transfected CHO cells with expression constructs encoding a PPAR-inducible firefly luciferase gene as well as a constitutively expressed renilla luciferase gene (transfection control). Transfection with these reporter constructs conferred PPAR-dependent expression of firefly luciferase to CHO cells as shown by the increase in firefly luciferase activity achieved by addition of the PAPRg agonist rosiglitazone to transfectants (Fig. 8A). Co-expression of sortilin and FABP7 in CHO-S/F cells resulted in a significantly higher relative activity of firefly luciferase as compared to CHO cells expressing either sortilin (CHO-S) or FABP7 (CHO-F) alone (Fig. 8B). PPAR-induced transcription of the firefly luciferase reporter in CHO-S/F cells was reduced by addition of exogenous apoE4 as compared with apoE3, substantiating the detrimental effects of apoE4 on the sortilin-dependent gene regulatory actions of FABP7 (Fig. 8C).

**Figure 8:**
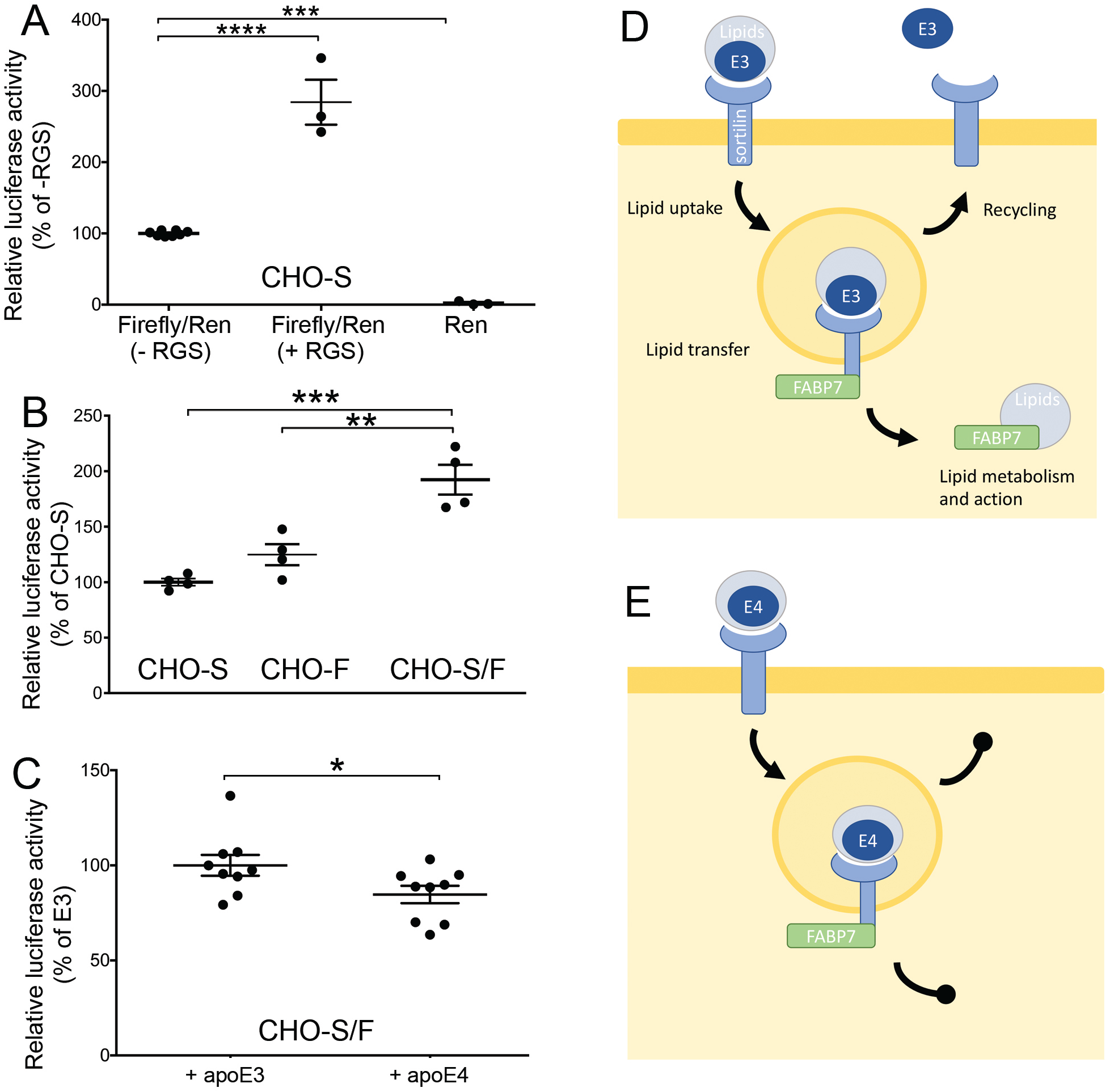
Interaction of sortilin and FABP7 in PPARg-dependent gene expression. (**A**) CHO cells stably expressing sortilin (CHO-S) were transfected with reporter gene constructs encoding a PPAR-responsive firefly luciferase gene (Luc) and a constitutively expressed renilla luciferase gene (Ren). Where indicated, transfectants were also treated with rosiglitazone (+ RGS) for 24 hours. Forty-eight hours after transfection, the activities of firefly and renilla luciferases were determined in cell lysates using a luminometer (n= 3-8 replicates per cell line). Values are given as ratio of firefly to renilla luciferase activity (mean ± SEM, condition without RGS as 100%). ***, p<0.001; ****, p<0.0001 (Student’s *t* test). (**B**) Replicate layers (n=4) of CHO cells stably expressing sortilin (CHO-S), FABP7 (CHO-F), or both proteins (CHO-S/F) were transfected with constructs encoding a PPAR-responsive firefly luciferase and a constitutively expressed renilla luciferase and analyzed for luciferase activity as described above. Values are given as ratio of firefly to renilla luciferase (mean ± SEM, CHO-S set to 100%). **, p<0.01; ***, p<0.001 (Student’s *t* test). (**C**) CHO-S/F were transfected with reporter gene constructs encoding a PPAR-responsive firefly luciferase reporter gene and a constitutively expressed renilla luciferase. Then, transfectants were treated with conditioned medium containing 5 μg/ml of human apoE3 or apoE4 for 24 hours. Forty-eight hours after transfection, the activities of firefly and renilla luciferases were determined in cell lysates using a luminometer (n= 9 replicates from 3 independent experiments per condition. Values are given as ratio of firefly to renilla luciferase (mean ± SEM, + apoE3 set to 100%). *, p<0.05 (Student’s *t* test). (**D-E**) Proposed model of the interaction of sortilin with FABP7 and apoE3 (D) or apoE4 (E) in cellular lipid metabolism (see discussion section for details).

## DISCUSSION

We propose a novel concept defining a role for sortilin in neuronal lipid metabolism (Fig. 8D-E). By binding of lipidated apoE from the extracellular space, sortilin enables neuronal uptake of essential lipids, including PUFA, the precursors to eCBs. Simultaneous binding of FABP7 to the cytoplasmic tail of the receptor facilitates spatial proximity of apoE and FABP7 in the early endocytic compartments and facilitates lipid transfer from the extracellular to the intracellular transport particle. In the presence of apoE3, proper recycling of sortilin and apoE3 to the cell surface releases lipid-laden FABP7, which directs further steps in intracellular metabolism and action of the lipids (Fig. 8D). By contrast, the known propensity of apoE4 to block recycling of sortilin (Asaro et al., 2020) impairs release of FABP7 and blocks subsequent steps in intracellular lipid handling and action (Fig. 8E).

FABP7 is a member of a class of chaperones that facilitate intracellular transport of lipids by increasing their solubility and rate of diffusion in the aqueous milieu of the cytosol (reviewed in (Matsumata et al., 2016; Moulle et al., 2012)). FABP7 shows highest affinity for docosahexaenoic acid (DHA), the most abundant ω3-PUFA in the brain, but also binds other fatty acid derivatives, including eCBs (Balendiran et al., 2000; Kaczocha et al., 2009; Xu et al., 1996). FABP7 directs cellular uptake (Kaczocha et al., 2009) and intracellular metabolism (Elmes et al., 2015) of fatty acids. In addition, it facilitates nuclear targeting of DHA and eCBs for activation of PPARs (Adida and Spener, 2006; Mita et al., 2010; Tripathi et al., 2017). Our data now document that FABP7 binds to sortilin and that this interaction controls stability and intracellular fate of the lipid carrier. Absence of sortilin alters subcellular distribution of FABP7 and decreases its half-life, an effect seen in CHO cells, in primary neurons, and in the brains of apoE3 mice deficient for sortilin. In recent transcriptomic analyses, no impact of apoE genotype on *Fabp7* transcript levels was reported in the murine brain or human cerebral organoids (Zhao et al., 2020a; Zhao et al., 2020b). These findings agree with our data that sortilin impacts post-translational mechanisms of FABP7 expression control (Figs. 4D and 6F). In cells, FABP7 localizes to the cytoplasm, plasma membrane and nucleus, in line with its role in lipid sorting in these compartments. Apart from a nuclear localization signal (Liang et al., 2006), little is known about the mechanisms that direct this chaperon to distinct cellular organelles. Our data document that FABP7 directly interacts with sortilin, as shown by coIP (Fig. 2A-B), and that it colocalizes with the receptor in endocytic and recycling compartments of the cell (Figs. 2D and 7F-I). At present, we can only speculate about the molecular mechanism whereby sortilin affects trafficking of FABP7. However, an activity for sortilin in intracellular sorting of lipid transport particles has been shown in hepatocytes, where this receptor affects secretion of apoB-containing lipoproteins (Kjolby et al., 2010; Sparks et al., 2016; Strong et al., 2012).

FABP7 is perceived as a protein predominantly expressed in specialized glia such as Bergmann glia in the cerebellum or ependyma in brain cortex (Kaczocha et al., 2009; Kurtz et al., 1994; Owada, 2008). Our studies show co-expression of FABP7 with sortilin in neurons in culture (Fig. 6A) and in murine and human brains (Fig. 5). Undoubtedly, neuronal expression is modest as compared to glia. Still, given the low abundance of FABP7-expressing glia in cortex, one may conclude that neurons likely contribute the bulk of FABP7 mass detected in lysates of murine and human brain cortex (Fig. 4). Why sortilin deficiency impacts stability of FABP7 in neurons (Fig. 6) but not astrocytes (Suppl. figure 1) is unclear at present, but lends further support for a molecular pathway unique to neuronal handling of PUFAs.

Loss of FABP7 activity in *Fabp7* gene-targeted mice results in disturbances in brain lipid homeostasis including a decrease in DHA content (Owada et al., 2006). A comparable phenotype is seen in *Sort1* KO mice that exhibit reduced levels of ω3-PUFA, DHA, as well as the eCB synaptamide (Asaro et al., 2020). As such changes are seen in the absence of any human APP transgenes in these mouse models, the effects of sortilin and apoE4 genotypes on brain lipid homeostasis are clearly not a secondary consequence of amyloidogenic processes. Whether *Fabp7* mutant mice are prone to developing AD-like pathologies has not been investigated thus far. However, low levels of DHA and eCBs are correlated with AD pathologies in patients and mouse models (Jung et al., 2012; Koppel et al., 2009; Maroof et al., 2014; Schaefer et al., 2006; Soderberg et al., 1991; Yuki et al., 2014); and so is low PPAR activity (Heneka et al., 2005; Pedersen et al., 2006). Jointly with the association of low FABP7 levels with the *APOEe4* genotype in human AD specimens shown by us, these data argue for the significance of a lipid transport machinery involving sortilin, apoE3, and FABP7 in brain health; and defects in this machinery as a primary risk factor of sporadic AD.

## MATERIALS AND METHODS

### Human specimens

For expression analysis, brain autopsy specimens from the frontal cortex of AD patients were obtained from the Netherlands Brain Bank (Netherlands Institute for Neuroscience, Amsterdam) and the MRC London Brain Bank for Neurodegenerative Diseases (Institute of Psychiatry, King’s College London). The ethnicity of samples was white. All material was collected from donors for or from whom a written informed consent for a brain autopsy and the use of the material and clinical information for research purposes had been obtained by the Netherlands Brain Bank or the MRC London Brain Bank. Detailed personal information, including age, gender, neuropathological stage, *APOE* genotype, of the individuals were provided by the brain banks (Additional file 1). Human post-mortem AD brain samples for immunohistochemistry were obtained from the archives of the Department of (Neuro) Pathology, Academic Medical Center (AMC), University of Amsterdam. Cases were selected from a retrospective searchable neuropathologic database, including cases with consent for post-mortem brain autopsy and use of their brain material and medical records for research (Additional file 2). The samples were used in compliance with the Declaration of Helsinki. The studies were approved by the Ethical Committees of the Academic Medical Center, Amsterdam (W11_073).

### Animal experimentation

The generation of *Sort1*^-/-^ (Jansen et al., 2007) and *Apoe* targeted replacement (Knouff et al., 1999) strains of mice has been described. The animals were kept on normal chow (4.5% crude fat, 39% carbohydrates). All animal experimentation was conducted in male mice on an inbred C57Bl6/J background following approval by local ethics committees (X9017/17). The animals were 11 to 13 weeks of age.

### Surface proteome analysis

Mixed hippocampal/cortical neuronal as well as astrocytic cultures were prepared from newborn WT or *Sort1* KO mice. The experimental details are given in the supplementary method section.

### Immunohistochemistry

For staining of human brain tissue, 3-4 μm paraffin sections were placed on poly-L-lysine coated slides and allowed to dry in an oven (37^0^C) overnight and then processed for immunohistochemistry described in detail elsewhere (Jesse et al., 2017). The sections were deparaffinized in xylene for 20 min and rehydrated in 100%, 96% and 75% ethanol for 3 min each followed by endogenous peroxidase quenching (0.3% H_2_O_2_ in methanol) for 20 min. For antigen retrieval, sections were heated in citrate buffer, pH 6 (DAKO), for 10 min in a pressure cooker. After washing in PBS, sections were incubated with primary antibody directed against FABP7 (kindly provided by T. Müller, MDC Berlin) at 4° C overnight. After washing in PBS, sections were incubated with secondary antibody (IL Immunologic, Duiven, The Netherlands) for 30 min at room temperature. DAB reagent (BrightDAB substrate kit; ImmunoLogic, Duiven, The Netherlands) was used to visualize antibody binding. The sections were then counter-stained with 6% hematoxylin for 3 min. All procedures were performed at room temperature.

For immunohistology of the murine brain, 10 μm coronal cryosections were incubated in blocking buffer (0.5% BSA, 10% donkey serum and 0.3% triton) for 1 hour. The sections were subsequently washed in PBS-T (1X PBS, 0.1% triton) for 10 minutes, followed by immunostaining with primary antibodies (diluted in blocking buffer) directed against NeuN (Abcam, #ab104224; 1:500), FABP7 (EMD Millipore, #ABN14; 1:200) and sortilin (R&D Systems, #AF2934; 1:200), followed by incubation with fluorophore-conjugated secondary antibodies (Alexa Fluor 488, 555, and 647 diluted 1:1000). The sections were washed, stained with DAPI (1:3000), and mounted onto glass slides for standard confocal microscopy.

### Expression analysis

Determinations of transcript and protein levels were performed by quantitative RT-PCR and SDS-PAGE, respectively, using standard protocols. Probes for transcript analyses were Taqman *Gapdh* (Mm99999915_g1), *Fabp7* (Mm00445225_m1), and *Gfap* (Mm01253033_m1). Fold change in transcript levels was calculated using the cycle threshold (CT) comparative method (2^-ddCT^) normalizing to CT values of internal control genes *Gapdh*. For semi-quantitative western blot analysis, the intensities of immunoreactive signals were quantified by densitometric scanning of blots using the Image Studio Lite software. Signal intensities were always normalized to loading controls detected on the same membrane. Primary antibodies used for immunodetection were directed against β-tubulin (abcam, ab6046; Millipore, CP06), FABP7 (Millipore, ABN14), FABP5 (Biozol Diagnostica, BVD-RD181060100-1), GAPDH (Genetex, GTX627408-01), apoE (Millipore, AB947), and sortilin (BD Transduction Laboratories^™^, 612101).

### PPAR-dependent luciferase assay

To investigate the transcriptional activity of PPARs, we used a peroxisome proliferator response element (PPRE)-driven luciferase reporter gene approach (Kim et al., 1998). In brief, CHO cell lines were transfected with a reporter construct containing three copies of the PPRE placed upstream of the thymidine kinase promoter-firefly luciferase fusion gene (PPRE X3-TK-luc, Addgene). In addition, the cells were transfected with a cytomegalovirus promoter-driven renilla luciferase reporter gene (pGL4.75 hRluc/CMV) used as transfection control. Where indicated, transfectants were treated with 5 μM of rosiglitazone (Sigma, R2408) or with conditioned medium containing 5 μg/ml of human apoE3 or apoE4 for 24 hours. Twenty-four to 48 hours after transfection, cells were lysed in buffer (50 mM Tris HCL, pH 7.4, 140 mM NaCl and 1% Triton X-100) and lysates used to assay luciferase activity in 96-well plates using the Dual-Luciferase Reporter Assay (Promega) according to the manufacturer’s instructions. The luminescence-based PPARs activity (firefly luciferase) was normalized to the internal renilla luciferase activity.

### Statistical analysis

For all *in vivo* experiments, an indicated number *n* is the number of mice per group used in an experiment. For primary culture experiments, an indicated number *n* is the number of independent neuronal or glial preparations (biological replicates) used for western blotting or qRT-PCR analyses. For co-localization studies in CHO cells, *n* is the number of cells analyzed in replicate experiments. Each mouse or human specimen, or biological replicate in a cell culture experiment represents a statistically independent experimental unit, which was treated accordingly as an independent value in the statistical analyses. Statistical analyses were performed using GraphPad Prism software. For data with two different conditions and/or time points, two-way ANOVA with Bonferroni multiple comparisons test was applied. To select differentially expressed proteins in the surface proteome analysis, a resampling ANOVA-based significance test was used. Further details of statistical analyses not included here are specified in the respective figure legends.

## Acknowledgment

We are indebted to T. Pasternack, K. Kampf, K. M. Pedersen, and C. Kruse for expert technical assistance. We also wish to thank H. Riezman (University of Geneva) for support and J. Bosset (Bioimaging Center UNIGE) for help with microscopy.

## Competing interests

The authors declare that they have no conflict of interest

## Funding

Studies were funded in part by the ERC (BeyOND No. 335692), the Helmholtz Association (AMPro), the Alzheimer Forschung Initiative (#18003), and the Novo Nordisk Foundation (NNF18OC0033928) to TEW; and by the Foundation for Polish Science co-financed by the European Union under the European Regional Development Fund (Homing program, POIR.04.04.00 00 5CEF/18 00) to ARM. Equipment for surface proteomics was provided in part by the Centre for Preclinical Research and Technology (CePT), co-sponsored by the European Regional Development Fund and Innovative Economy, The National Cohesion Strategy of Poland.

## Availability of data and materials

Data and materials are available from the corresponding author upon reasonable request.

## Ethics approval and consent to participate

Human brain samples were collected from donors for or from whom a written informed consent for a brain autopsy and the use of the material and clinical information for research purposes had been obtained by the Netherlands Brain Bank, the MRC London Brain Bank or the Department of (Neuro) Pathology, Academic Medical Center (AMC), University of Amsterdam. The latter samples were used in compliance with the Declaration of Helsinki. The studies were approved by the Ethical Committees of the Academic Medical Center, Amsterdam (W11_073).

## Animal experimentation

All animal experimentations were conducted in male mice on an inbred C57Bl6/J background following approval by local ethics committees (X9017/17).

## Authors’ contribution

AA, RS, OK, AC-S, MB, AR, ARM, and TR performed experiments and evaluated data. AA, MD, BK, EA, ARM, and TEW designed experiments and evaluated data. TEW wrote the manuscript. All authors read and approved the final manuscript.

## SUPPLEMENTARY METHODS

### Surface proteome analysis

Neurons were plated on poly-D-lysine coated plates (5×10^6^ cells per 10 cm plate) and maintained in Neurobasal medium (Invitrogen) supplemented with B27, glutamax, and penicillin/streptomycin (Invitrogen) and used for experiments at DIV10-12. Astrocytic cultures were prepared as published (Malik et al., 2020). Cells were plated on poly-L-lysine-coated flasks and cultured in DMEM with 10% FBS and penicillin/streptomycin. After 10 days *in vitro*, microglia was removed by shaking and the astrocytes were plated for experiments. Biotinylation and purification of surface proteins was performed using EZ-Link^™^ Sulfo-NHS-SS-Biotin (Thermo Fisher Scientific) and Neutravidin slurry (Pierce) as described (Malik et al., 2019). For mass spectrometry analyses, proteins were reduced with 5 mM TCEP for 60 min at 60°C. To block reduced cysteines, 200 mM MMTS at a final concentration of 10 mM was added and the sample was incubated at room temperature for 10 minutes. Proteins were digested overnight with 10 ng/μl trypsin and the resulting peptides were analyzed by LC-MS/MS for peptide identification and LC-MS for relative quantitation using a Q Exactive mass spectrometer (Thermo Scientific) coupled with a nanoAcquity LC system (Waters). To do so, the samples were desalted and concentrated on a nanoACQUITY UPLC Trapping Column (Waters). Further peptide separation was carried on a nanoACQUITY UPLC BEH C18 Column (Waters, 75 μm inner diameter; 250 mm long) with an acetonitrile gradient (5–35% over 160 minutes) in the presence of 0.1% formic acid at a flow rate of 250 nl/min. Qualitative LC-MS/MS measurements (i.e., peptide and protein identification) were carried out on samples pooled within each experimental group. Quantitative analyses of individual samples used separate survey scan LC-MS runs.

Pre-processed LC-MS/MS data files were submitted to the Mascot search engine (Matrix Science) and searched against *Mus musculus* protein entries from the SwissProt database using a target/decoy approach. The search parameters were as follows: enzyme: semitrypsin; number of missed cleavages: 1; ion mass error tolerances: ± 5 ppm (parent) and ± 0.01 Da (fragment); modifications: Methylthio C (fixed), Oxidation M (variable). The statistical significance of the identifications was assessed as described (Kall et al., 2008). Only peptide sequences with *q*-values ≤ 0.01 were considered as confidently identified.

Peptides identified in the LC-MS/MS analysis were quantified in individual LC-MS samples using a feature extraction procedure (Bakun et al., 2009). The calculated peptide abundances were log-transformed, normalized by a robust locally weighted regression smoother (LOESS) (Cleveland and Devlin, 1988), and rolled-up to relative protein abundances. To select differentially expressed proteins, a resampling ANOVA-based significance test was used.

### Cell culture experiments

To generate medium containing human apoE3 or apoE4, HEK293 cells were transiently transfected with expression constructs encoding for human apoE3 or apoE4. Cell supernatants were conditioned for 48 hours in medium without FCS (Chen et al., 2010). Alternatively, supernatants from primary astrocytes were conditioned for 48 hours in medium without FCS. The concentration of apoE in media batches from HEK293 or primary astrocytes was determined by western blot analysis comparing apoE signal intensities to those from serial dilutions of recombinant human apoE (MBL, JM-4699-500) included in the same blots.

For western blot and qRT-PCR analyses, neuronal or glial cultures were used at DIV 10-12. For immunocytochemistry, cells plated on coverslips were fixed with 4% PFA, incubated overnight at 4°C with primary antibodies directed against MAP2 (Synaptic systems, #188004; 1:250), FABP7 (EMD Millipore, #ABN14; 1:100) and sortilin (R&D Systems, #AF2934; 1:100), followed by incubation with fluorophore-labeled secondary antibodies (Alexa Fluor 488, 555, and 647 diluted 1:1000) and mounting onto glass slides using fluorescent mounting medium (Dako). Parental Chinese hamster ovary (CHO-K1) cells or cell clones stably overexpressing mouse sortilin and/or myc-tagged mouse FABP7 (Origene, MR200772) were grown in DMEM (Gibco) supplemented with 10% FCS (Gibco) and 1% penicillin/streptomycin (Gibco). Cell lysates for western blot analysis were generated using standard protocols.

### Protein interaction studies

Proximity ligation is an experimental approach to test the spatial proximity of proteins in cells. It employs oligonucleotide-conjugated antibodies directed against the target proteins, (here FABP7-myc and sortilin). Here, CHO-S/F cells stably overexpressing mouse sortilin and myc-tagged mouse FABP7 were grown on glass coverslips and treated for 24 hours with conditioned medium containing 5 μg/ml of human apoE3 or apoE4. Then, the cells were fixed with 4% PFA in PBS and subjected to proximity ligation assay (PLA) using orange PLA probes, followed by ligation and amplification performed according to the manufacturer’s recommendation (Sigma). Thereafter, the cells were incubated with primary and secondary antibodies to co-stain for the indicated marker proteins (anti-Rab5, Cell Signaling, #2143, 1:500; anti-Rab11, Cell Signaling, #5589, 1:500; and Alexa Fluor 647-conjugated secondary antibodies). Microscopy images were quantified using Fiji software. Colocalization analysis was performed with the Colocalization Threshold plugin with manual selection of the region of interest. Single z-plane images were used for calculating thresholded Mander’s coefficient (tM1). Analysis of protein localization by subcellular fractionation was carried out as previously described (Andersen et al., 2005).

## SUPPLEMENTARY TABLES

**Supplementary table 1:** List of selected proteins with altered cell surface localization in primary cultures of sortilin-deficient neurons or astrocytes

| ACC | Protein | Gene | ratio (KO/WT) | p-value |
| --- | --- | --- | --- | --- |
| **Neurons** |  |  |  |  |
| P15209 | BDNF/NT-3 growth factors receptor | *Trkb* | 0.724 | 0.020768 |
| Q01279 | Epidermal growth factor receptor | *Egfr* | 0.793 | 0.000107 |
| Q04519 | Sphingomyelin phosphodiesterase | *Asmase* | 1.53 | 0.005667 |
| Q8C0E2 | Vacuolar protein sorting-associated protein 26B | *Vps26b* | 1.56 | 0.0035004 |
| P51880 | Fatty acid-binding protein, brain (FABP7) | *Fabp7* | 1.52 | 0.033944 |
| P11404 | Fatty acid-binding protein, heart (FABP3) | *Fabp3* | 1.4 | 0.22835 |
| Q05816 | Fatty acid-binding protein, epidermal (FABP5) | *Fabp5* | 1.25 | 0.11495 |
| **Astrocytes** |  |  |  |  |
| P15209 | BDNF/NT-3 growth factors receptor | *Trkb* | 0.774 | 0.03436 |
| P28798 | Progranulin | *Grn* | 0.646 | 0.000474 |
| Q06335 | Amyloid-like protein 2 | *Aplp2* | 0.707 | 0.000877 |
| P41233 | ATP-binding cassette transporter ABCA1 | *Abca1* | 0.756 | 0.003095 |
| P51880 | Fatty acid-binding protein, brain (FABP7) | *Fabp7* | 0.94 | 0.61984 |
| P11404 | Fatty acid-binding protein, heart (FABP3) | *Fabp3* | not detected |  |
| Q05816 | Fatty acid-binding protein, epidermal (FABP5) | *Fabp5* | not detected |  |

ACC, Uniprot accession number; ratio (KO/WT), ratio between protein amounts in cell surface fractions from sortilin KO and WT primary cell cultures as determined by quantitative label-free proteomics; *p*-value is given based on a resampling ANOVA-based significance test.

**Supplementary table 2.** Patient samples examined by western blot analysis. Specimens were obtained from the Netherlands Brain Bank (NL; Netherlands Institute for Neuroscience, Amsterdam) and the MRC London Brain Bank for Neurodegenerative Diseases (UK; Institute of Psychiatry, King’s College London).

| Case No. | Age | Sex | Clinical diagnosis | Braak stage | ApoE Genotype | Brain Bank |
| --- | --- | --- | --- | --- | --- | --- |
| 94-110 | 82 | F | AD dementia | VI | 4/4 | NL |
| 94-037 | 63 | M | AD dementia | VI | 4/4 | NL |
| 94-082 | 64 | M | AD dementia | VI | 4/4 | NL |
| 98-060 | 66 | M | AD dementia | VI | 4/4 | NL |
| 94-091 | 83 | F | AD dementia | VI | 3/3 | NL |
| 94-025 | 85 | F | AD dementia | VI | 3/3 | NL |
| 94-016 | 86 | F | AD dementia | VI | 3/3 | NL |
| 93-140 | 87 | F | AD dementia | IV | 3/3 | NL |
| 97-009 | 89 | F | AD dementia | VI | 3/3 | NL |
| 92-022 | 91 | F | AD dementia | IV | 3/3 | NL |
| 93-011 | 92 | F | AD dementia | VI | 3/3 | NL |
| 99-114 | 92 | F | AD dementia | IV | 3/3 | NL |
| 99-123 | 93 | F | AD dementia | IV | 3/3 | NL |
| 96-020 | 58 | M | AD dementia | VI | 3/3 | NL |
| 94-028 | 70 | M | AD dementia | VI | 3/3 | NL |
| 95-077 | 72 | M | AD dementia | VI | 3/3 | NL |
| 90-066 | 89 | M | AD dementia | IV | 3/3 | NL |
| 92-024 | 89 | M | AD dementia | IV | 3/3 | NL |
| 99-064 | 89 | M | AD dementia | VI | 3/3 | NL |
| A197/88 | 81 | M | AD dementia | V-VI | 3/3 | UK |
| A005/96 | 89 | M | AD dementia | - | 3/3 | UK |
| A012/96 | 89 | M | AD dementia | V-VI | 3/3 | UK |
| A013/96 | 85 | M | AD dementia | V-VI | 3/3 | UK |
| A342/96 | 67 | M | AD dementia | V-VI | 3/3 | UK |
| A277/97 | 89 | M | AD dementia | V-VI | 3/3 | UK |
| A065/02 | 82 | M | AD dementia | V-VI | 3/3 | UK |
| A093/97 | 91 | M | AD dementia | V-VI | 4/4 | UK |
| A200/97 | 74 | M | AD dementia | - | 4/4 | UK |
| A213/94 | 81 | F | AD dementia | III-IV | 3/3 | UK |
| A335/94 | 72 | F | AD dementia | V-VI | 3/3 | UK |
| A125/95 | 88 | F | AD dementia | V-VI | 3/3 | UK |
| A044/96 | 84 | F | AD dementia | V-VI | 3/3 | UK |
| A097/96 | 90 | F | AD dementia | V-VI | 3/3 | UK |
| A291/96 | 60 | F | AD dementia | V-VI | 3/3 | UK |
| A407/96 | 75 | F | AD dementia | V-VI | 3/3 | UK |
| A025/97 | 80 | F | AD dementia | V-VI | 3/3 | UK |
| A026/98 | 80 | F | AD dementia | V-VI | 3/3 | UK |
| A028/98 | 70 | F | AD dementia | V-VI | 3/3 | UK |
| A207/98 | 91 | F | AD dementia | V-VI | 3/3 | UK |
| A133/99 | 92 | F | AD dementia | III-IV | 3/3 | UK |
| A053/95 | 76 | F | AD dementia | V-VI | 4/4 | UK |
| A240/95 | 75 | F | AD dementia | V-VI | 4/4 | UK |
| A366/95 | 79 | F | AD dementia | V-VI | 4/4 | UK |
| A073/96 | 89 | F | AD dementia | V-VI | 4/4 | UK |
| A133/97 | 92 | F | AD dementia | - | 4/4 | UK |
| A053/98 | 89 | F | AD dementia | V-VI | 4/4 | UK |

**Supplementary table 3.** Patients examined by immunohistology

| Case No. | Age | Sex | Post-mortem interval (hrs) | Clinical diagnosis | Thal Phase | Braak stage | CERAD neuritic plaque score | AD neuropathological change |
| --- | --- | --- | --- | --- | --- | --- | --- | --- |
| 1 | 71 | M | 12 | AD dementia | 4 | V-VI | 3 | A3, B3, C3 |
| 2 | 75 | M | 6 | AD dementia | 4 | V-VI | 2 | A3, B3, C2 |
| 3 | 82 | F | 24 | AD dementia | 4 | V-VI | 3 | A3, B3, C3 |

**Neuropathological assessments:** All cases were extensively characterized according to the routine protocol for neurodegenerative diseases. Neuropathological diagnosis of AD was made following the NIA-AA criteria including Thal phasing for A $\beta$ load, Braak-and- Braak- staging for NFTs, and CERAD neuritic plaque score to assess the density of neuritic plaques. (Clin Pathol. 2019 Nov;72(11):725-735;. Neuropathology of Neurodegenerative Diseases. (2014). In G. Kovacs (Ed.), Neuropathology of Neurodegenerative Diseases: A Practical Guide (pp. I-Ii). Cambridge: Cambridge University Press).

## SUPPLEMENTARY FIGURE LEGEND

**Supplementary figure S1.**
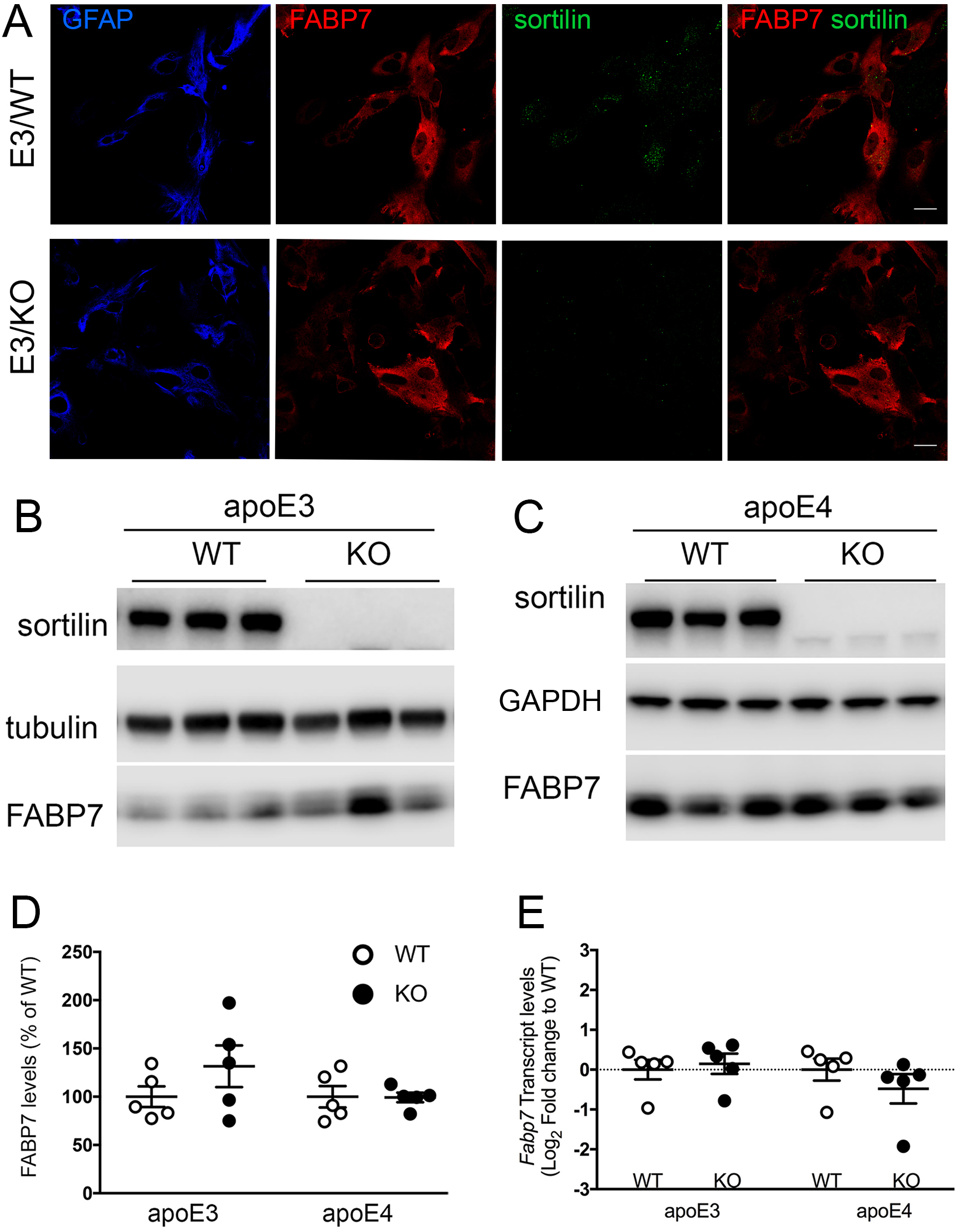
Sortilin deficiency does not impact FABP7 levels in primary astrocytes. (**A**) Immunodetection of FABP7 (red) and sortilin (green) in primary astrocytic cultures from apoE3 mice either (WT) or genetically deficient for *Sort1* (KO). FAPB7-stained cells are identified as astrocytes by expression of GFAP (blue). Merged images show co-expression of FABP7 and sortilin. Images represent single z-planes. Scale bar: 20 μm. (**B-C**) Exemplary western blot analyses of sortilin and FABP7 in primary astrocytes from apoE3 or apoE4 targeted replacement mice, either wild-type (WT) or homozygous for the *Sort1* null allele (KO). Detection of sortilin and tubulin served as controls. (**D**) Quantitative analysis of FABP7 levels in primary astrocyte cultures using densitometric scanning of replicate blots (exemplified in panels b and c). No genotype-dependent alterations in FABP7 levels were seen comparing cells of the indicated *APOE* and *Sort1* genotypes. Data are given as mean ± SEM with the respective WT level set to 100% (n=5 independent cultures per genotype). (**E**) Levels of *Fabp7* transcript were identical in primary astrocytes from apoE3- and apoE4-expressing mice, either WT or KO for *Sort1*, as determined by quantitative RT-PCR (n=5 independent cultures per genotype). Values are given as log2 fold change relative to the respective WT (set to value 0).

## Notes

### Competing Interest Statement

The authors have declared no competing interest.

